# Analysis of the DNA-binding domain of the *Pseudomonas aeruginosa* quorum sensing transcription factor LasR

**DOI:** 10.64898/2026.08.12.744480

**Authors:** Zichu Yang, Anvita Billa, Aditya S. Desai, Matthew R. Parsek, Ajai A. Dandekar

## Abstract

Many bacteria engage in quorum sensing (QS), a cell-cell communication system used to coordinate group behaviors. In one type of QS, acyl-homoserine lactone signals generated by LuxI homologs bind to LuxR homolog transcription factors, usually resulting in gene activation. The genome of *Pseudomonas aeruginosa* encodes three such LuxR homologs: LasR, RhlR, and QscR. Of these, LasR regulates the most genes, including that encoding RhlR. There is strong evidence that, during chronic infections, *lasR* and other genes encoding LuxR-type regulators are under strong selective pressure for mutations that both inactivate and modulate their function. Thus, we wondered if some mutations in the *lasR* gene might result in a protein with affinity for promoters usually regulated by the other LuxR homologs; to do so, we investigated the DNA-binding domain (DBD) of LasR through alanine substitution. As expected, we found that most alanine substitutions across the LasR DBD led to loss of function, as did previously identified clinical LasR DBD variants. Additionally, some alanine mutants were indistinguishable from the wild type. We describe a handful of variant LasR polypeptides that unexpectedly exhibit enhanced regulation on a RhlR-regulated gene, which conferred a fitness defect when competed against the wild type. Most other LasR variants had a competitive advantage. Our results suggest a pathway for expansion of the regulon of LuxR-homolog transcription factors, but also that such mutations may be disfavored due to the incurred metabolic burden.

**Importance:** Many bacteria generate chemical signals to alter gene expression in response to changes in population density, a phenomenon called quorum sensing. One type of quorum sensing relies on acyl-homoserine lactone (AHL) signals. In this type of quorum sensing, first described in the bioluminescent bacterium *Vibro fisheri*, a LuxI homolog produces the AHL, which binds to a LuxR homolog that typically activates gene expression. The opportunistic pathogen *Pseudomonas aeruginosa* has two such LuxR homologs, LasR and RhlR, each of which has its own specific regulon. We focused on the transcription factor LasR and investigated structural determinants of its binding to target promoters using an alanine substitution approach. We discovered that some DNA-binding mutations can expand the range of LasR-regulated genes. Our work provides insight into understanding what promoters LuxR homologs bind to and, more generally, how these proteins might evolve over time to change the group of genes that they regulate.

## Introduction

Quorum sensing (QS) is a widespread bacterial communication system that relies on the production of diffusible signals (1–3). One type of QS system, first characterized in the marine bacterium *Vibrio fischeri* (4, 5, 1), consists of a ligand-binding transcription factor, LuxR, and a cognate ligand synthase LuxI, whose expression is under the direct regulation of LuxR (5, 6). The activity of the transcription factor and the expression of the ligand synthase thus form an autoregulatory loop. LuxI and its homologs produce acyl-homoserine lactone (AHL) signals, with binding specificity related to the acyl group (1–3). The accumulation of AHL signals and transcriptional activation of LuxR-regulated genes results in coordinated, population-level gene expression.

One bacterium that engages in QS is *Pseudomonas aeruginosa,* an opportunistic pathogen of vulnerable populations including people who are immunocompromised and those with cystic fibrosis (CF) (7, 8). In the airways of people with these infections can be decades long and are associated with deterioration of lung function (9). Once established, *P. aeruginosa* infections can persist despite aggressive antibiotic treatment.

The *P. aeruginosa* genome encodes three complete QS systems (*las*, *rhl,* and *pqs*), (1–3). Two of these, Las and Rhl, are homologous to the *V. fischeri* system and involve synthesis of AHL signals by the synthases LasI and RhlI. LasI produces *N*-3-oxo-dodecanoyl-homoserine lactone (3OC12-HSL), which binds to and activates LasR. RhlI similarly produces *N*-butanoyl-homoserine lactone (C4-HSL) which binds to and activates RhlR. The *Pqs* signal, in contrast, is 2-heptyl-3-hydroxy-4-quinolone (“PQS”), which is synthesized by the proteins PqsABCD and PqsH. PQS binds to the receptor PqsR (also MvfR) (10). These three QS circuits have been described to be organized hierarchically: LasR activates expression of *rhlR* and *pqsR* (3), and each transcription factor has its own regulon (10–12). *P. aeruginosa* encodes an additional LuxR homolog gene, *qscR.* QscR does not have a paired AHL ligand synthase, and instead activates gene expression in the presence of several AHL signals, including 3OC12-HSL (13). QscR impairs LasR activity by an unresolved mechanism (14).

The *P. aeruginosa* QS hierarchy is adaptable, and variants that lack functional LasR polypeptides have been identified in numerous environments (15–19). Many of these isolates, unlike lab strains, engage in *rhl* QS independently of LasR (17, 18). There are also several naturally-occurring LasR variants that have partial function (17, 20).

The functional form of LasR is a symmetric homodimer. Each monomer consists of an N-terminal ligand-binding domain and a C-terminal DNA-binding domain (21). The LasR-3OC12-HSL interaction is important for dimerization and solubility, and induces LasR activity (11, 22, 23). This active form of LasR binds to target DNA sequences with unique motifs called Lux boxes (11, 3). Each of the LuxR homolog transcription factors in *P. aeruginosa* has a unique binding motif and distinct regulon (11, 24). In cases where a gene is regulated by both LasR and RhlR, the two transcription factors do not share the same binding motif but rather each has its own (11).

DNA-binding domains among LuxR homologs are structurally conserved with a classic helix-turn-helix motif. A previous analysis of the LuxR DNA-binding domain found that a majority of the residues within the motif are functionally important, either for DNA binding or RNA polymerase recruitment (6). The diversity of binding sequence motifs among LuxR homologs in *P. aeruginosa* inspires the question as to whether DNA-binding domain residues encode the binding specificity.

We reasoned that the hierarchy of QS circuits in *P. aeruginosa* allows us to ask two different and important questions: first, what are the important residues in the LasR polypeptide for activation of target promoters? and second, what is the potential for LuxR homologs to expand the suite of genes they activate through mutation? To do so we embarked on a systematic, alanine-scanning approach of the LasR DNA-binding domain (DBD). We characterized the QS activity of the resulting variants as well as a handful of previously identified DBD variants from clinical isolates (20). We investigated whether LasR mutants could potentially activate RhlR-regulated genes (11) and subsequently identified LasR DBD variants that partially activate the RhlR-specific *rhlA* promoter. These variants can confer a growth fitness disadvantage in strains that harbor them.

## Results

### A majority of alanine variants of LasR result in loss of function, but some retain function

LuxR homologs consist of a signal binding domain and a DNA-binding domain (3, 21). Because we are interested in which residues of the LasR DBD are involved with promoter binding, we constructed 50 alanine substitution mutants of LasR spanning amino acid residues 177 to 230 (Pfam PF00196), excluding 4 positions (residues 189, 206, 227 and 228) that are already alanine in the wild-type sequence. Our approach paralleled a previous study of the *V. fischeri* LuxR protein (6). We introduced these variant *lasR* alleles, along with the wild type (WT), into an episomal plasmid under the regulation of the arabinose-inducible P*_araBAD_* promoter and asked if they could activate gene expression. To do so, we generated a fusion of the LasR-regulated *rsaL* promoter (11, 20) with mScarlet. We used this *P_rsaL_-mScarlet* reporter, expressed on a secondary plasmid, and measured mScarlet fluorescence over growth. In all experiments we supplemented with 2 μM 3OC12-HSL and arabinose. To account for differences in timing of activation, we measured LasR activity by calculating the area under the curve in exponential phase (Figs. 1A and S1).

**Figure. 1.**
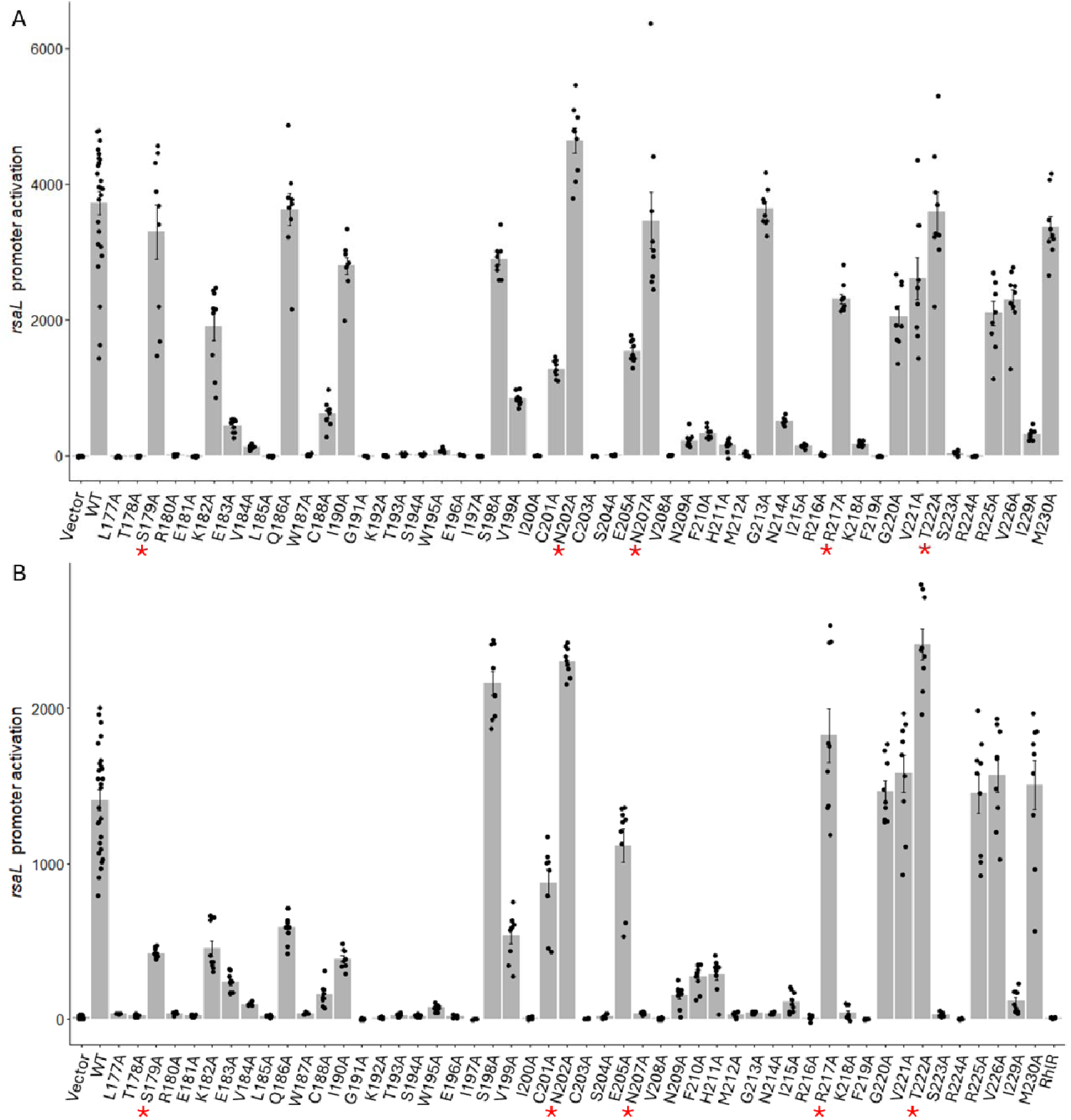
Activation of the LasR-regulated *rsaL* promoter by LasR DNA-binding domain variants. We measured activity by calculating the area under the curve after 14 h of growth using the P_rsaL_::mScarlet reporter in(A) Δ*lasR* background (B) Δ*lasR*Δ*rhlR*Δ*qscR* background. Asterisks indicate variants that were selected for further investigation.

We found that majority of the alanine substitution mutants were negative for LasR activity as measured by mScarlet fluorescence. However, of the 50 non-alanine residues, 8 substitutions did not alter LasR activity and 9 partially retained LasR activity. Next, to eliminate potential cross-activation by the other AHL responsive transcription factors, RhlR and QscR, we introduced the *P_rsaL_-mScarlet* reporter into PAO1Δ*lasR*Δ*rhlR*Δ*qscR* and repeated the experiment (Fig. 1B). We found that absence of RhlR and QscR did not affect the relative LasR activity to the WT for almost all of the LasR variants, apart from N207A and G213A, whose LasR activity was reduced in this background.

We then asked if absence of RhlR and QscR affected protein stability of these LasR variants in PAO1. We performed LasR western blots on lysates of cells expressing these variants in the PAO1Δ*lasR*Δ*rhlR*Δ*qscR* to determine if protein was present in the soluble fraction. Almost all did, with the notable exception of four (S177A, N207A, G213A and N214A) for which no band was apparent at the predicted LasR size (Fig. S1). Consistent with this apparent instability of the protein, we observed no LasR activity of these variants (Fig. 1A). Again, a notable variant was N207A, which only yielded a band by Western blotting at a size that corresponds to a LasR dimer.

### Some LasR variants appear to have expanded promoter targets

In addition to the ability to activate the LasR-specific *rsaL* promoter, we asked if these alanine substitution variants could also activate the *rhlA* promoter. *rhlA* is regulated exclusively by RhlR (11, 17, 25). We constructed a *P_rhlA_-mGreenlantern* fusion to assess the ability of these variants to activate this gene. To our surprise, we found multiple variants as well as the WT LasR had some *rhlA* promoter activity above background in this strain that lack RhlR (Fig. 2), although we suspect that this is a function of overexpression of LasR in this experiment. Nevertheless, one variant, N207A, appeared to have increased *rhlA* promoter activity (Fig. 2A).

**Figure 2.**
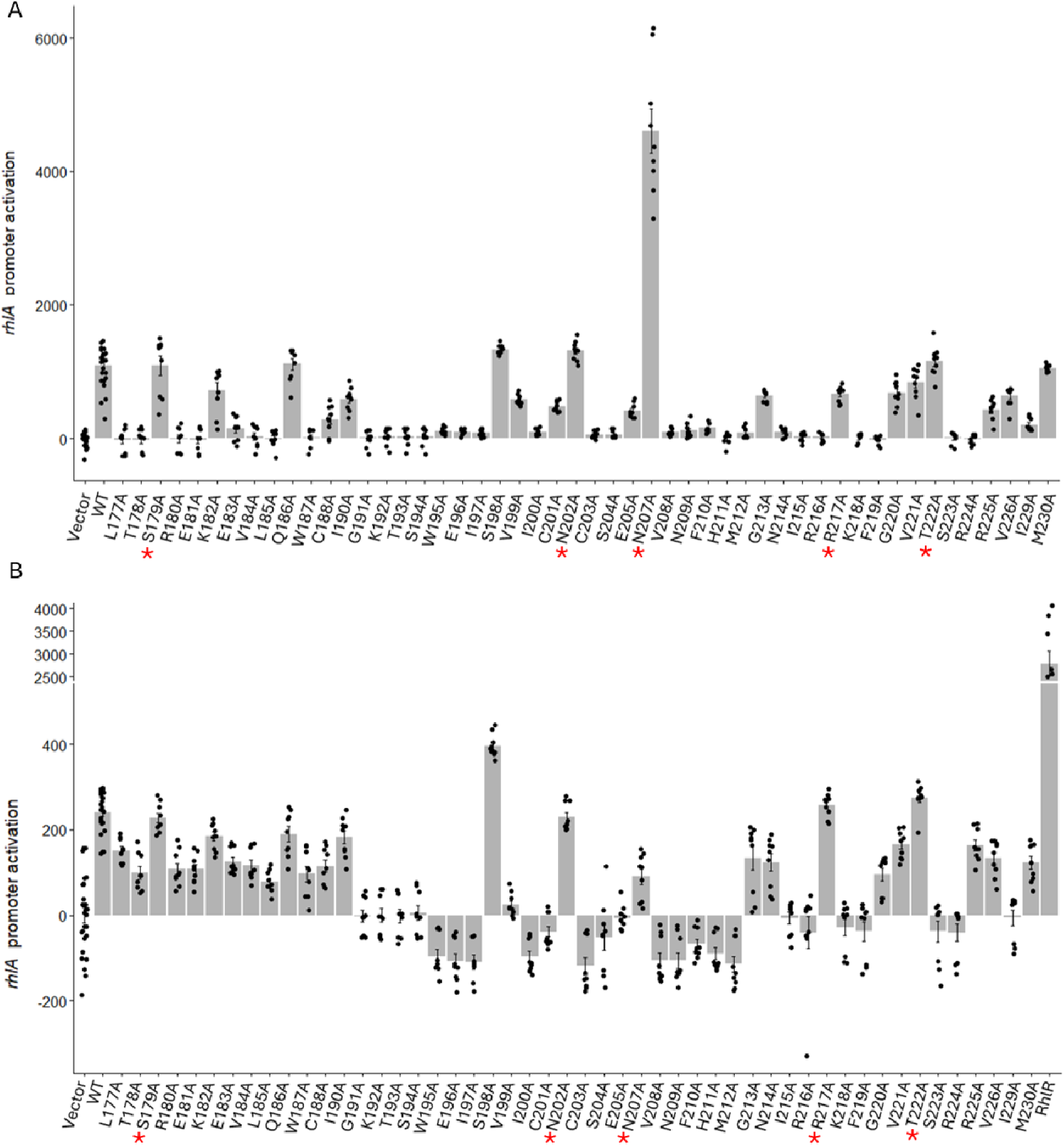
Activation of the RhlR-regulated *rhlA* promoter by LasR DNA-binding domain variants. We measured activity by calculating the area under the curve after 14 h of growth using the P_rhlA_::mGreenLantern reporter in (A) Δ*lasR* background (B) Δ*lasR*Δ*rhlR*Δ*qscR* background. Asterisks indicate variants that were selected for further investigation.

To test whether the *rhlA* activation was a consequence of the mutations or simply due to changes in LasR variant expression, we generated versions of LasR (S179A, N202A, N207A, R217A and T222A) through allelic exchange in PAO1 and PAO1 Δ*rhlR*Δ*qscR*. These strains express the variant LasR alleles from the native locus in the chromosome. We used the same promoter fusion reporters to examine *rsaL* and *rhlA* expression in these strains (Fig. 3). To our surprise, we found that all these variants, but not the control, displayed some *rhlA* expression, even without RhlR present in the genome (Fig. 3D). This result suggested that LasR can acquire, through mutation, some ability to regulate the *rhlA* promoter. The highest reporter activity was observed for N207A, but even this variant required RhlR to be present for full expression of the *rhlA* promoter (Fig. 3C and D).

**Figure 3.**
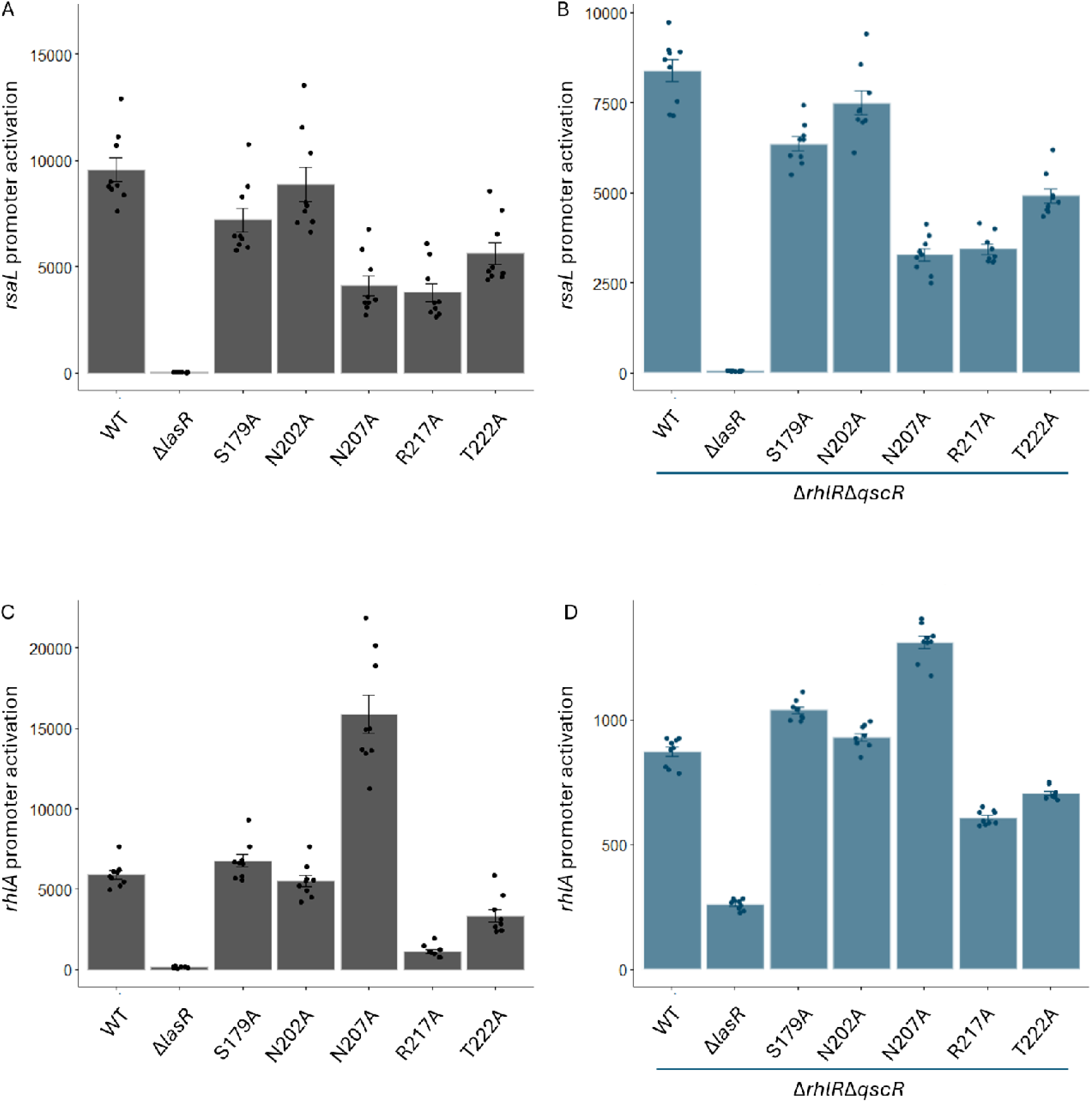
QS-regulated promoter activity from chromosomal expression of selected LasR variants. (A and B) *las* QS activity measured by area under the curve at 14 h using the P_rsaL_::mScarlet reporter in (A) Δ*lasR* background (B) Δ*lasR*Δ*rhlR*Δ*qscR*; (C and D) RhlR dependent promoter activity measured by area under the curve at 14 h using the P_rhlA_::mGreenLantern reporter in (C) Δ*lasR* background (D) Δ*lasR*Δ*rhlR*Δ*qscR* background.

We next asked whether this apparent activation of P_RhlA_ by LasR results in production of products traditionally thought to be only Rhl-regulated. Therefore, we tested these variants for their ability to produce rhamnolipids and pyocyanin (26). PAO1 strains expressing the LasR variants demonstrated enhanced rhamnolipid and pyocyanin production compared to WT in the presence of wild-type *rhlR*, but did not in the Δ*rhlR*Δ*qscR* background (Fig. 4). This result suggests that apparent promoter cross-activation that we observed does not compensate fully for the lack of RhlR.

**Figure 4.**
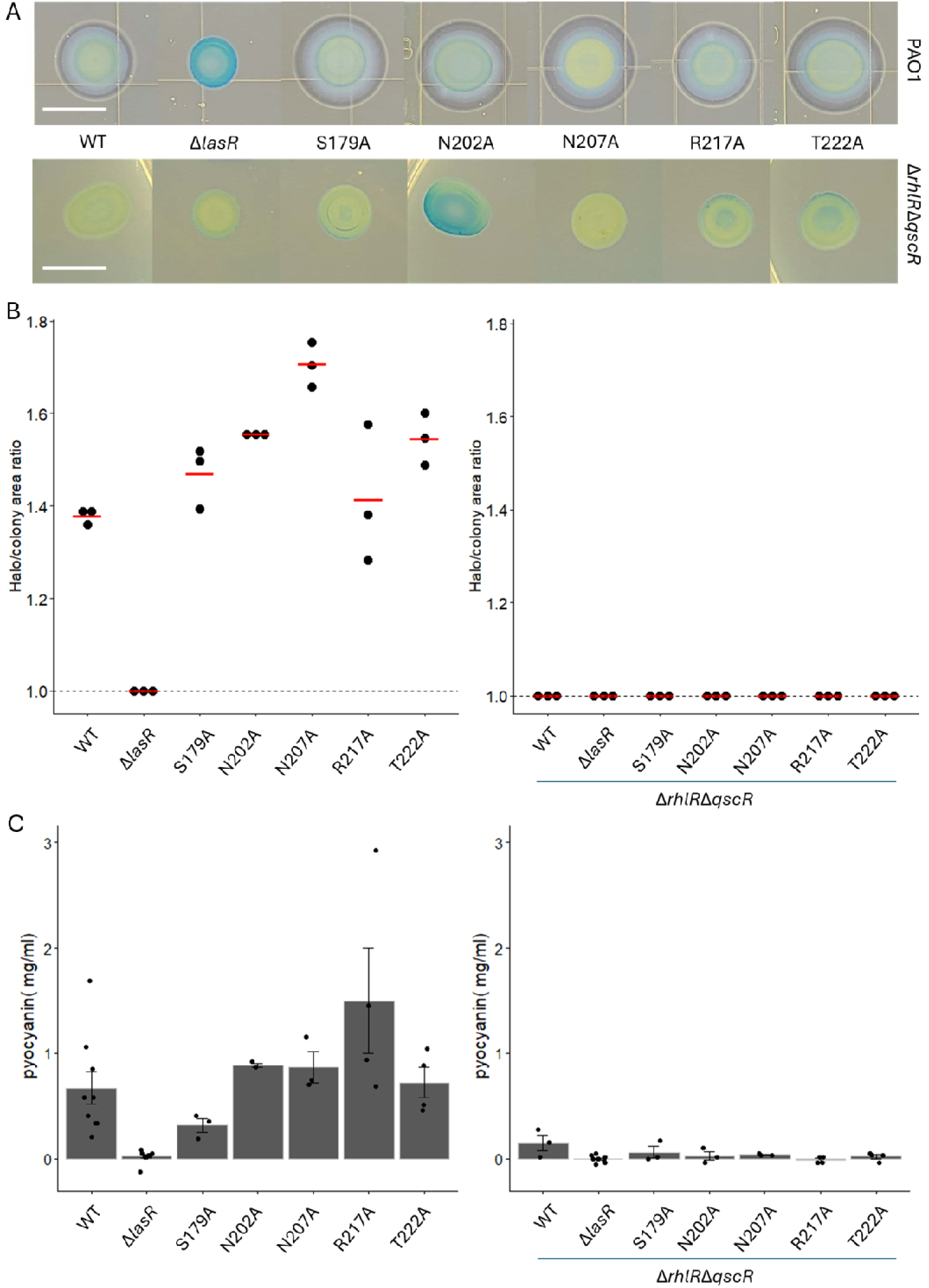
RhlR is necessary for LasR variants to exhibit rhamnolipid production and pyocyanin production from (A) Rhamnolipid detection plates from overnight cultures at 37°C followed by 72 hr incubation at room temperature. Scale bar is 10mm. (B) Quantification of colony size and clearing halo formation on rhamnolipid detection plates. (C) Pyocyanin production measured from stationary culture in buffered LB at OD_600_ of 3.0

### Characterization of naturally-occurring LasR DBD variants

In addition to the alanine substitution variants, we also examined LasR DBD variants (R216W, R216W-M230I, V226I, and A231V) that we recently identified as the most common amongst CF clinical isolates in the pseudomonas.com database (20, 27). We first asked if these variants could produce stable LasR variant proteins and assessed the protein stability of these natural variants with arabinose-induced overexpression in PAO1Δ*lasR*Δ*rhlR*Δ*qscR* as described above. All four of the natural variants expressed the LasR peptide at the expected size (Fig. S1). With the knowledge that these natural variants encoded stable protein, we next asked if they could activate gene expression. Among the three non-alanine residues previously identified in LasR DBD variants from clinical isolates, R216 is the least permissive to alanine substitution while V226 and M230 were only moderately affected (Fig. 1). We thus hypothesized that R216W would be null and the V226I variant would retain *las* activity. To best characterize these variants, we produced their chromosomal versions of (R216W, R216W-M230I, V226I, and A231V) through allelic exchange and monitored their impact on *rsaL* and *rhlA* promoter activity, as described above. LasR R216W did not activate either the *rsaL* or *rhlA* promoter, phenocopying the R216A variant (Fig. 5). The presence of a R216W-M230I double mutant in several unrelated isolates made us wonder if the M230I substitution was compensatory. However, in the conditions of our experiment, it was not: this variant phenocopied a Δ*lasR* mutant. The same was true for the V226I and A231V variants (Fig. 5).

**Figure 5.**
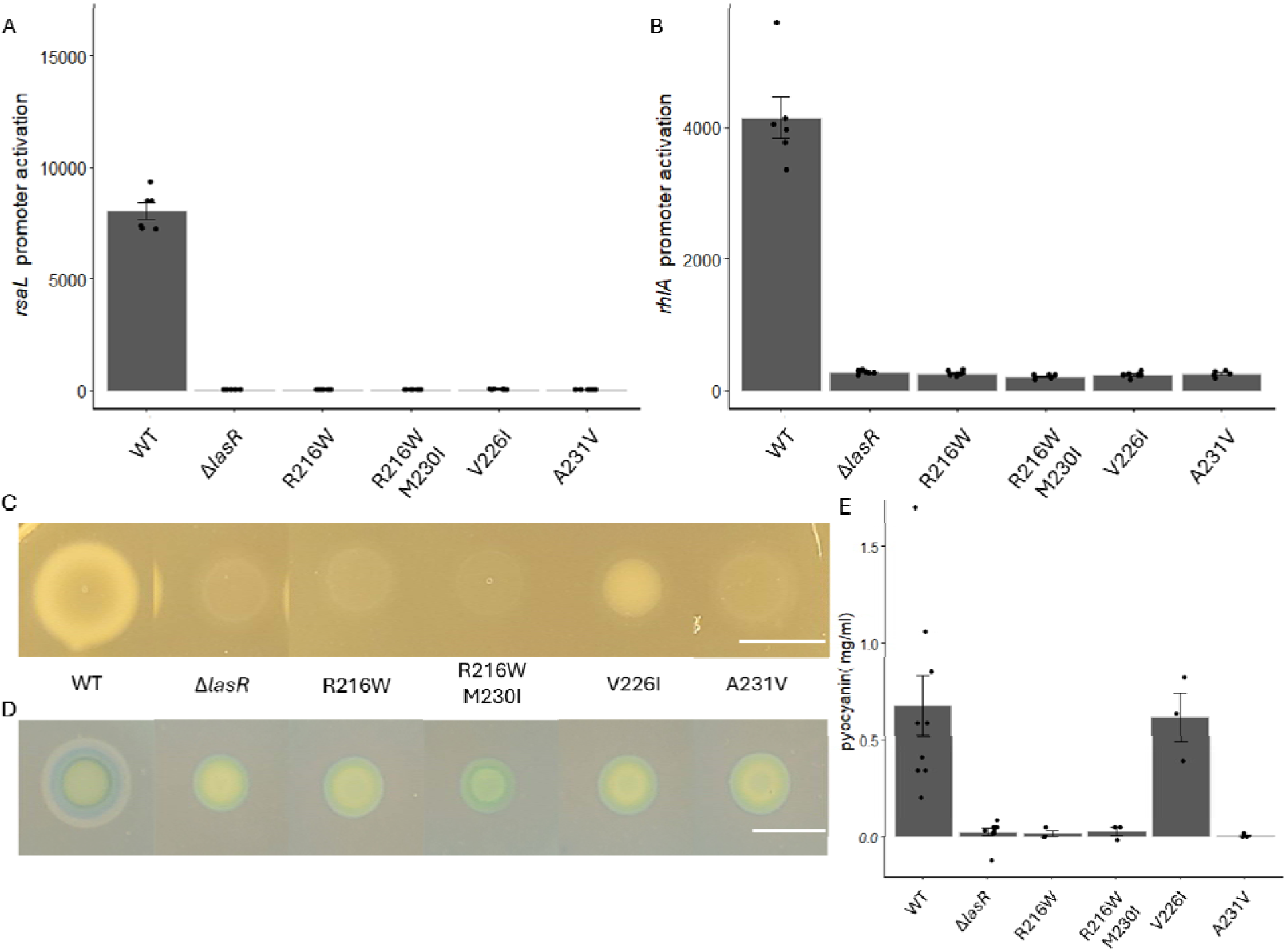
Characterization of LasR DNA-binding-domain natural variants. (A) *las* QS activity measured by area under the curve at 14 h using the P_rsaL_::mScarlet reporter in the Δ*lasR* background. (B) RhlR dependent promoter activity measured by area under the curve at 14 h using the P_rhlA_::mGreenLantern reporter in the Δ*lasR* background. (C) Representative colony growth and proteolysis phenotypes on casein agar plates. LasR V226I was the only variant with visible colony growth. Scale bar is 10mm. (D) Representative rhamnolipid production. None of the natural variants formed a halo. Scale bar is 10mm. (E) Pyocyanin production measured from stationary culture in buffered LB at OD_600_ of 3.0.

We validated these *rsaL* promoter activity results by growing *P. aeruginosa* on a minimal medium containing casein as the sole carbon and energy source (“casein broth”), which requires it to activate QS to obtain carbon and energy (15, 20). *P. aeruginosa* uses the LasR-and RhlR-regulated exoprotease elastase to break down casein into amino acids and peptides that can be used as carbon and energy sources. On casein-infused agar plates, we found the R216W, R216W-M230I, and A231V variants all phenocopied Δ*lasR* with no visible biomass buildup as expected while V226I had some level of growth, albeit reduced relative to WT and with no visible protease production (Fig. 5C). We next measured rhamnolipid and pyocyanin production by strains harboring these variants, as described above (Fig. 5D and E). All variants produced rhamnolipid at the same level as a Δ*lasR* mutant strain. R216W, R216W-M230I, and A231V again phenocopied Δ*lasR* in pyocyanin production, while V226I exhibited nearly wild-type levels of pyocyanin production.

Together, these results suggested that among the four variants, V226I uniquely retained some level of QS activity. Because this *las* activity did not correspond with our *rsaL* promoter activity assays, we speculate V226I has reduced regulon compared to WT, which is consistent with its diminished rhamnolipid production and a prior report from our group (20).

### LasR variant induction of *rhlA* correlates with a competitive disadvantage against the WT

The results so far suggest that a handful of LasR variants may expand the range of genes they activate, while most do not. We recently reported that the V226I variant, discussed in the prior section, has a competitive advantage over both the WT and LasR-null strains (20). We asked whether other mutations, particularly the ones that appear to partially activate *rhlA*, would have an advantage or (more likely) a disadvantage in competition with the WT.

We first assessed our collection of alanine substitution variants in both PAO1 and PAO1 Δ*rhlR*Δ*qscR* for their ability to individually grow on casein agar plates (Fig. 6A). Ten variants (S179A, N202A, N207A, R217A, T222A, in both backgrounds) were able to develop measurable biomass on casein plates. Except for R217A and the LasR-null control, all variants in the PAO1 background formed a halo of protease production around the colony, while only N207A formed such a halo in the PAO1Δ*rhlR*Δ*qscR* background. Because both LasR and RhlR regulate elastase production, we reasoned this result is consistent with N207A having enhanced transcriptional activity of the corresponding promoters in both PAO1 and PAO1 Δ*rhlR*Δ*qscR* (Fig. 3C and D).

**Figure 6.**
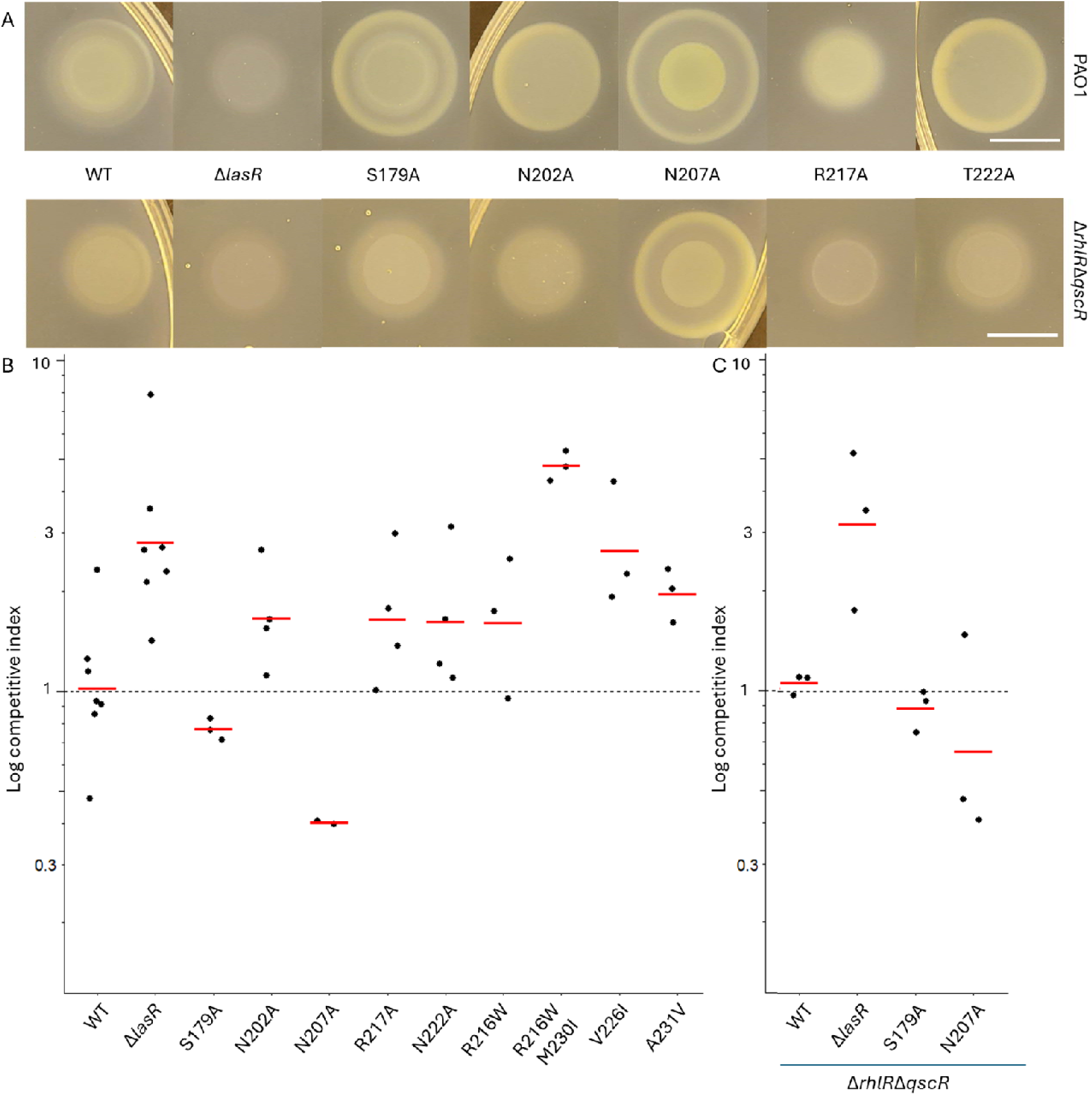
LasR variants that appear to activate *rhlA* confer a competitive disadvantage in casein broth. (A) Representative colony growth and proteolysis phenotypes on casein agar plates. Scale bar is 10mm. (B and C) Competitive index (final frequency / initial frequency) of individual LasR variants after 18 h competition with the WT. Starting ratio was at 1 (variant):10 (WT). Background for the variants: (B) PAO1; (C) PAO1Δ*rhlR*Δ*qscR*. Red bars indicate the mean of three independent experiments.

To test LasR variant strain fitness, we grew these in competition with the WT in casein broth. LasR-null mutants have a growth fitness advantage over the WT because they are relieved of the metabolic burden of QS while the WT incurs the cost of QS activation and elastase production for the whole population (18, 20, 28, 29) .

We measured the fitness all the LasR variants, in both the PAO1 and Δ*rhlR*Δ*qscR* backgrounds, against PAO1. We inoculated each individually at a 1:10 starting ratio (variant : WT). All naturally-occurring variants outcompeted the WT, as did most of the alanine-substitution variants, consistent with the idea that these mutations were inactivation. However, the S179A and N207A variants, which retained some *rhlA* promoter activity, were outcompeted by the WT (Fig. 6B). We hypothesized that this impaired fitness was due to an added cost burden attributable to the LasR variant itself or somehow due to activation of RhlR. To ask this question, we competed the S179A and N207A variants in the Δ*rhlR*Δ*qscR* against wild-type PAO1.These strains retained a competitive disadvantage against the WT (Fig. 6C), suggesting that the fitness disadvantages are due to the LasR variants themselves.

## Discussion

We are interested in the process by which LuxR homologs might evolve over time to regulate more or fewer genes. To gain insight into this question, we focused on the *P. aeruginosa* LasR DBD, generating a collection of variants, either through alanine scanning or by recreating mutant alleles observed in clinical isolates (20, 27). We reasoned that we would gain insight into residues that are important for binding to target DNA, and those that are permissive to substitution. One feature of *P. aeruginosa* QS that makes it particularly suited to this kind of analysis is the presence of genes encoding two additional LuxR homologs, RhlR and QscR, providing a kind of internal control for LuxR-regulated gene expression, as these two transcription factors activate a different suite of genes than LasR.

Unsurprisingly, we found that most substitutions in the LasR DBD led to loss of function. We also identified several residues that modulated LasR activity in an unexpected manner, including enhanced regulation of *rhlA*, which is typically regulated by RhlR alone (11). Additionally, three out of four clinical isolate DBD variants result in loss of function, while the previously-reported V226I variant (20) had significantly reduced activity.

Our promoter reporter fusion assay results are consistent with the previous analysis of the DNA-binding domain of LuxR (6), which demonstrated that the majority of the residues in the helix-turn-helix domain are important for its function. Our reporter fusion assays similarly identified two clustered regions (191 to 197, and 208 to 213) that show little tolerance to alanine substitutions, coalescing with the two annotated helices (194 to 201, and 205 to 220) of the HTH motif (Fig. 1). This pattern of clustering non-permissive residues also exists in LuxR but only within the second helix of the HTH motif. Only four residues among the HTH motif were reported as strongly reducing activity in LuxR while 16 residues strongly reduced LasR activity within the same region (Fig. 1). In addition, we confirmed that 7 amino acids that are conserved among LuxR homologs -- G191, K192, I197, I200, V208, N214 and K218 -- are essential for LasR activity (6). These comparisons indicate LasR DBD is less permissive to alanine substitutions than LuxR. While it is unclear as to why this might be the case, one possibility is that the stringency results from the need to have more DNA binding specificity to avoid cross-activation from the other LuxR homologs present in *P.aeruginosa*.

We were surprised that a handful of alanine substitutions in the DBD led to RhlR-independent *rhlA* promoter activation (Fig. 3D). This effect was detected only at the level of transcription (not production of rhamnolipid), but one intriguing possibility is that these LasR variants have broadened or altered their binding affinity as compared to the WT. Another possible interpretation is that LasR somehow indirectly regulates *rhlA* in the absence of RhlR, although there is no evidence to support this idea. Nevertheless, these LasR variants do appear to increase QS activity overall, demonstrated in their increase in rhamnolipid and pyocyanin production (Fig 4). The N207A variant stands out amongst this group: it uniquely boosts *rhlA* expression much more in the presence of RhlR than without, suggesting this variant might have a mechanism of regulation of the *rhl* regulon involving interaction with RhlR, including, potentially, heightened *rhlR* transcription.

In addition to investigating their impact of LasR on target promoter activity, we also wondered if binding affinity mutations might exhibit altered fitness in liquid culture. Using an established method to measure competitive fitness (18, 20) we found that individuals harboring LasR variants that exhibited enhanced *rhlA* promoter regulation had a fitness disadvantage as compared to the WT (Fig. 6). There was an interesting variant among the naturally-occurring LasR mutants, V226I. We and others have previously observed emergence of this variant in multiple laboratory evolution experiments and in clinical settings (20). Unlike other variants, this V226I appears to reduce the metabolic cost of QS by reducing rhamnolipid and elastase production while maintaining pyocyanin production (Fig. 5). Overall, these results are consistent with the concept that addition of genes to the quorum regulon of any LuxR homolog would require a specific selective pressure to be advantageous; otherwise, it is likely simply to increase the metabolic burden of QS.

Our work provides a framework for understanding both the structural determinants of LasR binding to target DNA and, more broadly, understanding how LuxR homologs might evolve over time to change the type and number of genes that they regulate. This process might occur iteratively with mutations in the signal synthase (30) to result in new signal-receptor pairs, or it might occur with no change in signal specificity as bacteria adapt to new environmental niches. Future work might focus on understanding what genes LasR variants regulate, how promoter regulation differs between LuxR homologs, and more generally understanding what DNA sequences are bound to by LuxR DNA-binding domains.

## Materials and Methods

### Bacterial strains and growth conditions

Bacterial strains and plasmids used in this study are listed in Supplemental tables S1 and S2. *E. coli* was grown in lysogeny broth (LB) at 37°C with shaking (250 rpm). *P. aeruginosa* was grown in LB buffered with 50 mM 3-(N-morpholino) propane sulfonic acid, (MOPS, pH 7.0) (buffered LB) or a minimal medium called photosynthesis medium (31) with 1% (wt/vol) casein sodium salt (casein broth), both at 37°C with shaking (250 rpm). For plasmid maintenance and selection in *P. aeruginosa*, the following antibiotics were used: carbenicillin, 200 μg/ml, gentamicin 50 μg/ml (maintenance) or 100 μg/ml (selection). In *E. coli*, ampicillin was added 100 μg/ml or gentamicin at 10 μg/ml.

### Construction of the LasR DBD variant plasmid library and chromosomal variants

LasR DBD variant fragments were ordered as synthetic DNA from Twist Bioscience and subsequently cloned into the pJN(Ap) plasmid backbone (32) between EcoRI and SacI sites with NEBuilder Hifi DNA assembly master mix (NEB). Assembled plasmids were transformed into and then maintained in DH5α cells (Invitrogen). All plasmids were confirmed via whole plasmid sequencing (Plasmidsaurus).

To create chromosomal LasR variants we used a homologous recombination-based two-step allelic exchange with sucrose counterselection approach, as previously described (18). We cloned the variant *lasR* sequence into the previously constructed pEXG2 lasR knockout plasmid with NEBuilder Hifi DNA assembly master mix (NEB) to create the cognate knock-in constructs, which were then electroporated into either PAO1 Δ*lasR* or PAO1 Δ*lasR*Δ*rhlR*Δ*qscR* backgrounds. All chromosomal variants were confirmed by whole genome sequencing (Plasmidsaurus).

### Reporter activity assays

A dual reporter plasmid pIY126 was constructed by three-part Gibson assembly via NEBuilder Hifi DNA assembly master mix (NEB) with P_rsaL_::mScarlet and P_rhlA_::mGreenLantern in addition to linearized XZ385 backbone (33). We used the same promoter regions for P_rsaL_ and P_rhlA_ as previously described (20). Both promoter activities were measured using a Biotek Synergy H1 microplate reader, as described in previous studies (18, 34). Briefly, colonies of strains with the transcriptional reporter pIY126 were inoculated into LB-MOPS supplemented with Gm50 and incubated at 37°C with shaking overnight. Cultures were back-diluted to an OD_600_ of 0.01 and grown until OD_600_ of 0.1. Exponential phase cultures were used to inoculate wells in a 96-well plate to an OD_600_ of 0.01 with a 200 μl final volume. Plates were incubated at 37°C with shaking for 14 hours in the microplate reader. Green fluorescence (excitation 489 nm, emission 520 nm), red fluorescence (excitation 570 nm, emission 605 nm) and OD_600_ were measured every 15 minutes. Where appropriate, 3OC12-HSL (Cayman) was added to a final concentration of 2 μM and arabinose to a final concentration 0.05%), at the time of inoculation.

### Immunoblotting

We assessed the relative levels of soluble LasR in the PAO1 Δ*lasR*Δ*rhlR*Δ*qscR* background with in-trans expression of arabinose inducible LasR variants using published methods (32, 35). Briefly, overnight cultures in LB-MOPS were diluted 1:100 in fresh LB-MOPS and then grown to OD_600_ of 0.4,at which point 0.1% arabinose and 2 μM of 3OC12-HSL (Cayman) were added to induce LasR expression and stabilize the protein. Induced cultures were then grown to OD_600_ of 2.0. Cells were then pelleted by centrifugation at 4°C and suspended in LasR purification buffer (25 mM Tris-HCl pH 7.8, 150 mM NaCl, 1 mM ethylenediaminetetraacetic acid, 1 mM dithiothreitol, 0.5% Tween-20, 10% glycerol, 2 µM 3OC12-HSL). We sonicated the suspensions and centrifuged lysates at 26,000 rpm for 20 minutes at 4°C, collected the supernatant and quantified protein concentrations by NanoDrop. Normalized samples were separated by SDS-PAGE and subsequently transferred to a PVDF membrane. After overnight blocking in TBST-milk (0.1% Tween 20, 5% skim milk), the membrane was treated with polyclonal antibodies against LasR (Covance; 1:1000 dilution) and protein detection was with a secondary goat anti-rabbit antibody (IRdye 800CW).

### Casein agar protease assessment

Casein agar was produced as previously described (36). We spotted 5 µL overnight cultures in LB-MOPS at OD_600_ of 4.0 were casein agar plates. Colony growth and clearing halo from casein proteolysis were captured after overnight growth at 37°C.

### Rhamnolipid measurement

Relative rhamnolipid production levels were quantified using a previously described method (34). Briefly, cultures of *P. aeruginosa* were grown in LB-MOPS overnight to OD_600_ of 4.0 at 37°C with shaking. Of this culture, 20 µL was spotted onto methylene blue-containing rhamnolipid detection plates (0.6% [wt/vol] Na_2_HPO_4_, 0.3% [wt/vol] KH_2_PO_4_, 0.05% [wt/vol] NaCl, 1.5% [wt/vol] Noble agar, 0.1 mM CaCl_2_, 2 mM MgSO_4_, 0.2% glucose, 0.05% glutamate, 0.0005% methylene blue, 0.02% cetyltrimethylammonium bromide, and 0.05% casamino acids). These plates were then incubated at 37°C overnight and subsequently at room temperature in dark for 96 hours. Each colony was measured for the colony and halo area (mm^2^) and we ascertained the ratio of the halo formation to colony size.

### Pyocyanin measurement

Cultures of *P. aeruginosa* were grown in LB-MOPS overnight at 37°C with shaking and pyocyanin was extracted from 4 ml culture supernatant with 4ml chloroform, and then extracted from the chloroform phase with an equal volume of 0.2 M HCl. Final pyocyanin concentration (mg/ml) was quantified by multiplying measured absorbance at 520 nm by 17.072 (37).

### Casein broth competition assay

We conducted competitions between mCherry-tagged PAO1 LasR variant strains and GFP-tagged wild-type PAO1. Three biological replicates of each strain were grown overnight in buffered LB and normalized to OD_600_ of 4.0. 150 µl of the WT overnight culture was mixed with 15 ul of the competing strain culture to form the inoculum at starting ratio 1:10, which was then added to 3 ml of casein broth. At the start of experiment and at the 18-hour timepoint,10 µl samples of the casein culture were washed in fresh PM casein and diluted 1:500 in PBS and the sample population was enumerated by a BD Accuri C6 flow cytometer, as previously described (20).

## Acknowledgements

This work was funded in part by NIH grants R35GM152107 (to AAD) and R01AI077628 and R01AI183692 (to MRP). We thank Andrew Frando, Nicole Smalley, and Amy Schaefer for technical assistance.

## References

1. Fuqua WC, Winans SC, Greenberg EP. 1994. Quorum sensing in bacteria: the LuxR-LuxI family of cell density-responsive transcriptional regulators. J Bacteriol 176:269–275.

2. Waters CM, Bassler BL. 2005. Quorum sensing: cell-to-cell communication in bacteria. Annu Rev Cell Dev Biol 21:319–346.

3. Miranda SW, Asfahl KL, Dandekar AA, Greenberg EP. 2022. *Pseudomonas aeruginosa* quorum sensing. Adv Exp Med Biol 1386:95–115.

4. Engebrecht J, Nealson K, Silverman M. 1983. Bacterial bioluminescence: isolation and genetic analysis of functions from *Vibrio fischeri*. Cell 32:773–781.

5. Engebrecht J, Silverman M. 1984. Identification of genes and gene products necessary for bacterial bioluminescence. Proc Natl Acad Sci U S A 81:4154–4158.

6. Egland KA, Greenberg EP. 2001. Quorum sensing in *Vibrio fischeri*: analysis of the LuxR DNA binding region by alanine-scanning mutagenesis. J Bacteriol 183:382–386.

7. Alhede M, Bjarnsholt T, Givskov M, Alhede M. 2014. *Pseudomonas aeruginosa* biofilms: mechanisms of immune evasion. Adv Appl Microbiol 86:1–40.

8. Malhotra S, Hayes D, Wozniak DJ. 2019. Cystic fibrosis and *Pseudomonas aeruginosa*: the host-microbe Interface. Clin Microbiol Rev 32:e00138–18.

9. Rosenfeld M, Faino AV, Qu P, Onchiri FM, Blue EE, Collaco JM, Gordon WW, Szczesniak R, Zhou Y-H, Bamshad MJ, Gibson RL. 2023. Association of *Pseudomonas aeruginosa* infection stage with lung function trajectory in children with cystic fibrosis. Journal of Cystic Fibrosis 22:857–863.

10. Xiao G, He J, Rahme LG. 2006. Mutation analysis of the *Pseudomonas aeruginosa mvfR* and *pqsABCDE* gene promoters demonstrates complex quorum-sensing circuitry. Microbiology (Reading) 152:1679–1686.

11. Schuster M, Urbanowski ML, Greenberg EP. 2004. Promoter specificity in *Pseudomonas aeruginosa* quorum sensing revealed by DNA binding of purified LasR. Proc Natl Acad Sci U S A 101:15833–15839.

12. Soto-Aceves MP, Smalley NE, Schaefer AL, Greenberg EP. The relationship between *pqs* gene expression and acylhomoserine lactone signaling in *Pseudomonas aeruginosa*. J Bacteriol 206:e00138–24.

13. Fuqua C. 2006. The QscR quorum-sensing regulon of *Pseudomonas aeruginosa*: an orphan claims its identity. J Bacteriol 188:3169–3171.

14. Ding F, Oinuma K-I, Smalley NE, Schaefer AL, Hamwy O, Greenberg EP, Dandekar AA. 2018. The *Pseudomonas aeruginosa* orphan quorum sensing signal receptor QscR regulates global quorum sensing gene expression by activating a single linked operon. mBio 9:10.1128/mbio.01274-18.

15. Sandoz KM, Mitzimberg SM, Schuster M. 2007. Social cheating in *Pseudomonas aeruginosa* quorum sensing. Proc Natl Acad Sci U S A 104:15876–15881.

16. Hoffman LR, Kulasekara HD, Emerson J, Houston LS, Burns JL, Ramsey BW, Miller SI. 2009. *Pseudomonas aeruginosa lasR* mutants are associated with cystic fibrosis lung disease progression. J Cyst Fibros 8:66–70.

17. Feltner JB, Wolter DJ, Pope CE, Groleau M-C, Smalley NE, Greenberg EP, Mayer-Hamblett N, Burns J, Déziel E, Hoffman LR, Dandekar AA. 2016. LasR variant cystic fibrosis isolates reveal an adaptable quorum-sensing hierarchy in *Pseudomonas aeruginosa*. mBio 7:e01513–16.

18. Kostylev M, Kim DY, Smalley NE, Salukhe I, Greenberg EP, Dandekar AA. 2019. Evolution of the *Pseudomonas aeruginosa* quorum-sensing hierarchy. Proceedings of the National Academy of Sciences 116:7027–7032.

19. Scribner MR, Stephens AC, Huong JL, Richardson AR, Cooper VS. The nutritional environment is sufficient to select coexisting biofilm and quorum sensing mutants of *Pseudomonas aeruginosa*. J Bacteriol 204:e00444–21.

20. Bellamoroso KR, Kostylev M, Smalley NE, Greenberg EP, Dandekar AA. 2026. Inactivation of MexT in *Pseudomonas aeruginosa* PAO1 destabilizes cooperation and favors the emergence of a unique quorum sensing variant. Journal of Bacteriology 208:e00434–25.

21. Bottomley MJ, Muraglia E, Bazzo R, Carfì A. 2007. Molecular insights into quorum sensing in the human pathogen *Pseudomonas aeruginosa* from the structure of the virulence regulator LasR bound to its autoinducer. Journal of Biological Chemistry 282:13592–13600.

22. Urbanowski ML, Lostroh CP, Greenberg EP. 2004. Reversible acyl-homoserine lactone binding to purified *Vibrio fischeri* LuxR protein. J Bacteriol 186:631–637.

23. Churchill MEA, Chen L. 2011. Structural basis of acyl-homoserine lactone-dependent signaling. Chem Rev 111:68–85.

24. Gilbert KB, Kim TH, Gupta R, Greenberg EP, Schuster M. 2009. Global position analysis of the *Pseudomonas aeruginosa* quorum-sensing transcription factor LasR. Mol Microbiol 73:1072–1085.

25. Keegan NR, Colón Torres NJ, Stringer AM, Prager LI, Brockley MW, McManaman CL, Wade JT, Paczkowski JE. 2023. Promoter selectivity of the RhlR quorum-sensing transcription factor receptor in *Pseudomonas aeruginosa* is coordinated by distinct and overlapping dependencies on C4-homoserine lactone and PqsE. PLoS Genet 19:e1010900.

26. Pearson JP, Pesci EC, Iglewski BH. 1997. Roles of *Pseudomonas aeruginosa las* and *rhl* quorum-sensing systems in control of elastase and rhamnolipid biosynthesis genes. J Bacteriol 179:5756–5767.

27. Winsor GL, Griffiths EJ, Lo R, Dhillon BK, Shay JA, Brinkman FSL. 2016. Enhanced annotations and features for comparing thousands of *Pseudomonas* genomes in the *Pseudomonas* genome database. Nucleic Acids Res 44:D646–653.

28. Wang M, Schaefer AL, Dandekar AA, Greenberg EP. 2015. Quorum sensing and policing of *Pseudomonas aeruginosa* social cheaters. Proc Natl Acad Sci USA 112:2187–2191.

29. Smalley NE, An D, Parsek MR, Chandler JR, Dandekar AA. 2015. Quorum sensing protects *Pseudomonas aeruginosa* against cheating by other species in a laboratory coculture model. Journal of Bacteriology 197:3154–3159.

30. Eldar A. 2011. Social conflict drives the evolutionary divergence of quorum sensing. Proc Natl Acad Sci U S A 108:13635–13640.

31. Kim M-K, Harwood CS. 1991. Regulation of benzoate-CoA ligase in *Rhodopseudomonas palustris*. FEMS Microbiol Lett 83:199–203.

32. Wellington Miranda S, Cong Q, Schaefer AL, MacLeod EK, Zimenko A, Baker D, Greenberg EP. 2021. A covariation analysis reveals elements of selectivity in quorum sensing systems. eLife 10:e69169.

33. Zheng X, Gomez-Rivas EJ, Lamont SI, Daneshjoo K, Shieh A, Wozniak DJ, Parsek MR. 2024. The surface interface and swimming motility influence surface-sensing responses in *Pseudomonas aeruginosa*. Proceedings of the National Academy of Sciences 121:e2411981121.

34. Frando A, Parsek RS, Omar J, Groleau M-C, Trottier MC, Smalley NE, Déziel E, Dandekar AA. 2025. Modulation of the *Pseudomonas aeruginosa* quorum sensing cascade by MexT-regulated factors. mBio 16:e02941–25.

35. Schuster M, Greenberg EP. 2007. Early activation of quorum sensing in *Pseudomonas aeruginosa* reveals the architecture of a complex regulon. BMC Genomics 8:287.

36. Chen R, Déziel E, Groleau M-C, Schaefer AL, Greenberg EP. 2019. Social cheating in a Pseudomonas aeruginosa quorum-sensing variant. Proc Natl Acad Sci USA 116:7021– 7026.

37. Kurachi M. 1958. Studies on the biosynthesis of pyocyanine. (II) : Isolation and determination of pyocyanine. Bulletin of the Institute for Chemical Research, Kyoto University.

