## Supplementary figure and tables for "Analysis of the DNA-binding domain of the *Pseudomonas aeruginosa* quorum sensing transcription factor LasR"

**Figure S1.** Immunoblots of LasR variant peptides expressed from an episomal plasmid in the PAO1 *ΔlasRΔrhlRΔqscR* background.

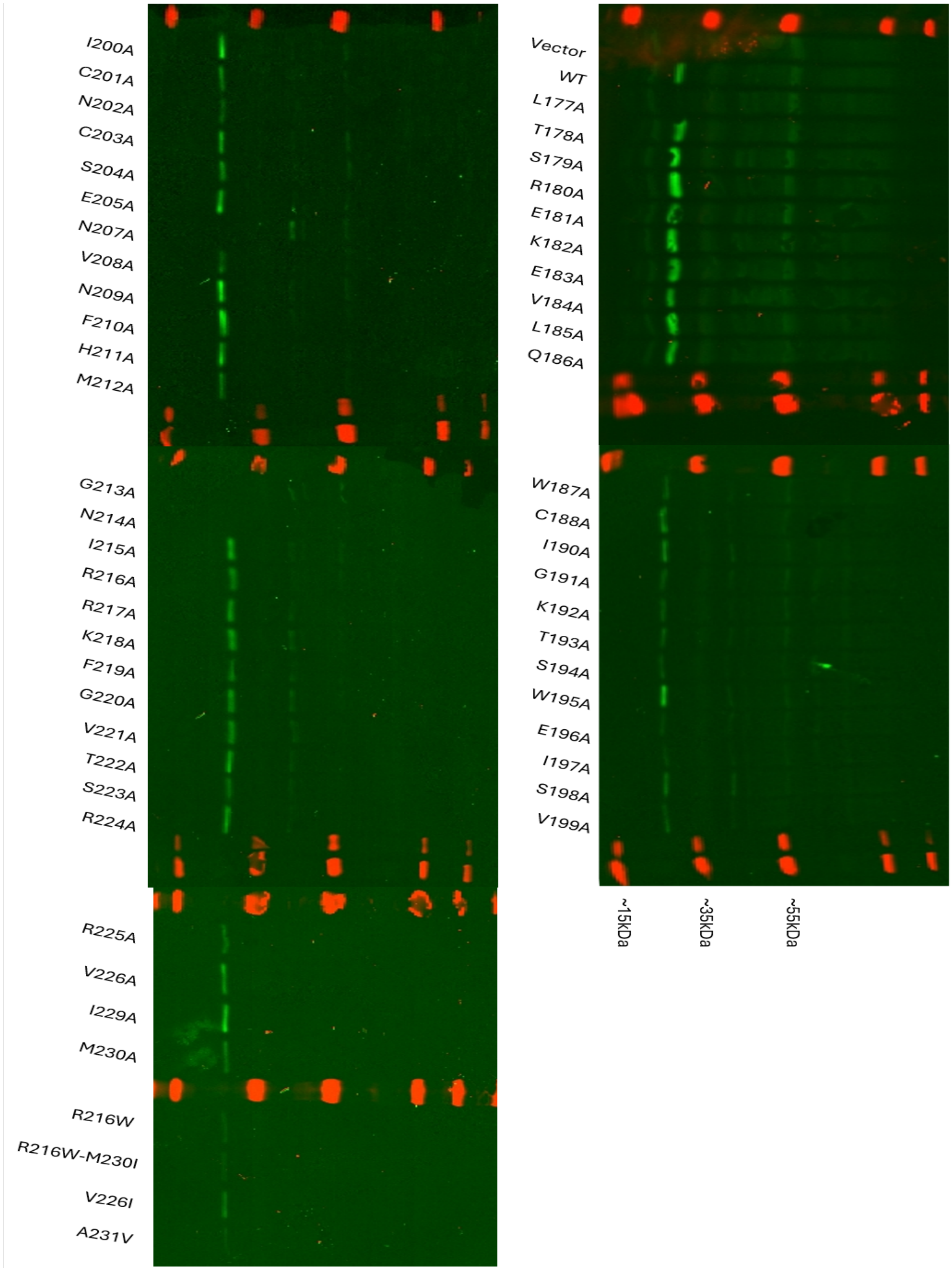

**Table S1.** Strains used in this study

| Strain Name | Description | Source |
|---|---|---|
| ***P. aeruginosa*** | | |
| mPAO1 | Wild-type *Pseudomonas aeruginosa* (NCBI accession number NZ_CP027857) | (1) |
| PAO1 Δ*lasR* | PAO1 Δ*lasR* | (2) |
| PAO1 Δ*lasR*Δ*rhlR*Δ*qscR* | PAO1 Δ*lasR*Δ*rhlR*Δ*qscR* | (3) |
| PAO1 Δ*rhlR*Δ*qscR* | PAO1 Δ*rhlR*Δ*qscR* with wild-type lasR | This work |
| PAO1 LasR S179A | PAO1 natively expressing LasR S179A variant* | This work |
| PAO1 Δ*rhlR*Δ*qscR* LasR S179A | PAO1 Δ*rhlR*Δ*qscR* natively expressing LasR S179A variant | This work |
| PAO1 LasR N202A | PAO1 natively expressing LasR N202A variant | This work |
| PAO1 Δ*rhlR*Δ*qscR* LasR N202A | PAO1 Δ*rhlR*Δ*qscR* natively expressing LasR N202A variant | This work |
| PAO1 LasR N207A | PAO1 natively expressing LasR N207A variant | This work |
| PAO1 Δ*rhlR*Δ*qscR* LasR N207A | PAO1 Δ*rhlR*Δ*qscR* natively expressing LasR N207A variant | This work |
| PAO1 LasR R217A | PAO1 natively expressing LasR R217A variant | This work |
| PAO1 Δ*rhlR*Δ*qscR* LasR R217A | PAO1 Δ*rhlR*Δ*qscR* natively expressing LasR R217A variant | This work |
| PAO1 LasR T222A | PAO1 natively expressing LasR T222A variant | This work |
| PAO1 Δ*rhlR*Δ*qscR* LasR T222A | PAO1 Δ*rhlR*Δ*qscR* natively expressing LasR T222A variant | This work |
| PAO1 LasR R216W | PAO1 natively expressing LasR R216W variant | This work |
| PAO1 LasR R216W-M230I | PAO1 natively expressing LasR R216W-M230I variant | This work |
| PAO1 LasR V226I | PAO1 natively expressing LasR V226I variant | (4) |
| PAO1 LasR A231V | PAO1 natively expressing LasR A231V variant | This work |
| PAO1-Tn7-gfp | mPAO1 with constitutively expressed *gfp* at Tn7 site | (5) |
| PAO1-Tn7-mCherry | mPAO1 with constitutively expressed *mCherry* at Tn7 site | (6) |
| PAO1 Δ*lasR*-Tn7-mCherry | PAO1 Δ*lasR* with constitutively expressed *mCherry* at Tn7 site | This work |
| PAO1 Δ*lasR*Δ*rhlR*Δ*qscR*-Tn7-mCherry | PAO1 Δ*lasR*Δ*rhlR*Δ*qscR* with constitutively expressed *mCherry* at Tn7 site | This work |
| PAO1 Δ*rhlR*Δ*qscR*-Tn7-gfp | PAO1 Δ*rhlR*Δ*qscR* with wild-type *lasR* with constitutively expressed *gfp* at Tn7 site | This work |
| PAO1 Δ*rhlR*Δ*qscR*-Tn7-mCherry | PAO1 Δ*rhlR*Δ*qscR* with wild-type *lasR* with constitutively expressed *mCherry* at Tn7 site | This work |
| PAO1 LasR S179A-Tn7-mCherry | PAO1 natively expressing LasR S179A variant and constitutively expressing mCherry at Tn7 site | This work |
| PAO1 Δ*rhlR*Δ*qscR* LasR S179A-Tn7-mCherry | PAO1 Δ*rhlR*Δ*qscR* natively expressing LasR S179A variant and constitutively expressing *mCherry* at Tn7 site | This work |
| PAO1 LasR N202A-Tn7-mCherry | PAO1 natively expressing LasR N202A variant and constitutively expressing *mCherry* at Tn7 site | This work |

| | | |
|---|---|---|
| PAO1 Δ*rhlR*Δ*qscR* LasR N202A-Tn7-mCherry | PAO1 Δ*rhlR*Δ*qscR* natively expressing LasR N202A variant and constitutively expressing *mCherry* at Tn7 site | This work |
| PAO1 LasR N207A-Tn7-mCherry | PAO1 natively expressing LasR N207A variant and constitutively expressing *mCherry* at Tn7 site | This work |
| PAO1 Δ*rhlR*Δ*qscR* LasR N207A-Tn7-mCherry | PAO1 Δ*rhlR*Δ*qscR* natively expressing LasR N207A variant and constitutively expressing *mCherry* at Tn7 site | This work |
| PAO1 LasR R217A-Tn7-mCherry | PAO1 natively expressing LasR R217A variant and constitutively expressing *mCherry* at Tn7 site | This work |
| PAO1 Δ*rhlR*Δ*qscR* LasR R217A-Tn7-mCherry | PAO1 Δ*rhlR*Δ*qscR* natively expressing LasR R217A variant and constitutively expressing *mCherry* at Tn7 site | This work |
| PAO1 LasR T222A-Tn7-mCherry | PAO1 natively expressing LasR T222A variant and constitutively expressing *mCherry* at Tn7 site | This work |
| PAO1 Δ*rhlR*Δ*qscR* LasR T222A-Tn7-mCherry | PAO1 Δ*rhlR*Δ*qscR* natively expressing LasR T222A variant and constitutively expressing *mCherry* at Tn7 site | This work |
| PAO1 LasR R216W-Tn7-mCherry | PAO1 natively expressing LasR R216W variant and constitutively expressing *mCherry* at Tn7 site | This work |
| PAO1 LasR R216W-M230I-Tn7-mCherry | PAO1 natively expressing LasR R216W-M230I variant and constitutively expressing *mCherry* at Tn7 site | This work |
| PAO1 LasR V226I-Tn7-mCherry | PAO1 natively expressing LasR V226I variant and constitutively expressing *mCherry* at Tn7 site | This work |
| PAO1 LasR A231V-Tn7-mCherry | PAO1 natively expressing LasR A231V variant and constitutively expressing *mCherry* at Tn7 site | This work |
| ***E. coli*** | | |
| NEB DH5α | fhuA2 Δ(argF-lacZ)U169 phoA glnV44 Φ80 Δ(lacZ)M15 gyrA96 recA1 relA1 endA1 thi-1 hsdR17 | NEB |
| S17-1 λ pir | recA pro hsdR RP4-2Tc::Mu-Km::Tn7 | (7) |

*Additional detail about the variants is provided in Table S2.

**Table S2.** Plasmids used in this study

| Plasmid | Description | Source |
|---|---|---|
| pIY126 | pBBR1MCS5-PrsaL-mScarlet-tet2-PrhlA-mGL-t0t1, generated based on XZ385 (8) | This work |
| pJN(Ap) | pJN105(Ampicillin resistant version) | (9) |
| pIY63/ pJN105LasR-L177A-Amp | pJN105 based expression vector containing LasR with CT to GC nucleotide substitution at position 529 and 530, resulting in L177A amino acid substitution | This work |
| pIY64/ pJN105LasR-E181A-Amp | pJN105 based expression vector containing LasR with A to C nucleotide substitution at position 542, resulting in E181A amino acid substitution | This work |
| pIY65/ pJN105LasR-R224A-Amp | pJN105 based expression vector containing LasR with CGC to GCG nucleotide substitution at position 670 to 672, resulting in R224A amino acid substitution | This work |
| pIY66/ pJN105LasR-T178A-Amp | pJN105 based expression vector containing LasR with A to G nucleotide substitution at position 532, resulting in T178A amino acid substitution | This work |
| pIY67/ pJN105LasR-R180A-Amp | pJN105 based expression vector containing LasR with CGG to GCC nucleotide substitution at position 538 to 540, resulting in R180A amino acid substitution | This work |
| pIY68/ pJN105LasR-L185A-Amp | pJN105 based expression vector containing LasR with TT to GC nucleotide substitution at position 553 and 554, resulting in K182A amino acid substitution | This work |
| pIY69/ pJN105LasR-G191A-Amp | pJN105 based expression vector containing LasR with G to C nucleotide substitution at position 572, resulting in G191A amino acid substitution | This work |
| pIY70/ pJN105LasR-I197A-Amp | pJN105 based expression vector containing LasR with AT to GC nucleotide substitution at position 589 and 590, resulting in I197A amino acid substitution | This work |
| pIY71/ pJN105LasR-V208A-Amp | pJN105 based expression vector containing LasR with AA to GC nucleotide substitution at position 544 and 545, resulting in K182A amino acid substitution | This work |
| pIY72/ pJN105LasR-H211A-Amp | pJN105 based expression vector containing LasR with CAT to GCA nucleotide substitution at position 631 to 633, resulting in H211A amino acid substitution | This work |
| pIY73/ pJN105LasR-N214A-Amp | pJN105 based expression vector containing LasR with AA to GC nucleotide substitution at position 640 and 641, resulting in N214A amino acid substitution | This work |

| | | |
|---|---|---|
| pIY74/ pJN105LasR-K218A-Amp | pJN105 based expression vector containing LasR with AAG to GCA nucleotide substitution at position 652 to 654, resulting in K218A amino acid substitution | This work |
| pIY75/ pJN105LasR-S223A-Amp | pJN105 based expression vector containing LasR with T to G nucleotide substitution at position 666, resulting in S223A amino acid substitution | This work |
| pIY76/ pJN105LasR-S179A-Amp | pJN105 based expression vector containing LasR with AGC to GCG nucleotide substitution at position 535 to 537, resulting in S179A amino acid substitution | This work |
| pIY77/ pJN105LasR-K182A-Amp | pJN105 based expression vector containing LasR with AA to GC nucleotide substitution at position 544 and 545, resulting in K182A amino acid substitution | This work |
| pIY78/ pJN105LasR-E183A-Amp | pJN105 based expression vector containing LasR with AA to CC nucleotide substitution at position 548 and 549, resulting in E183A amino acid substitution | This work |
| pIY79/ pJN105LasR-V184A-Amp | pJN105 based expression vector containing LasR with T to C nucleotide substitution at position 551, resulting in V184A amino acid substitution | This work |
| pIY80/ pJN105LasR-Q186A-Amp | pJN105 based expression vector containing LasR with CA to GC nucleotide substitution at position 556 and 557, resulting in Q186A amino acid substitution | This work |
| pIY81 pJN105LasR-W187A-Amp | pJN105 based expression vector containing LasR with TG to GC nucleotide substitution at position 559 and 560, resulting in W187A amino acid substitution | This work |
| pIY82/ pJN105LasR-S194A-Amp | pJN105 based expression vector containing LasR with AG to GC nucleotide substitution at position 580 and 581, resulting in S194A amino acid substitution | This work |
| pIY83/ pJN105LasR-W195A-Amp | pJN105 based expression vector containing LasR with TG to GC nucleotide substitution at position 583 and 584, resulting in W195A amino acid substitution | This work |
| pIY84/ pJN105LasR-F210A-Amp | pJN105 based expression vector containing LasR with TT to GC nucleotide substitution at position 628 and 629, resulting in F210A amino acid substitution | This work |
| pIY85/ pJN105LasR-T222A-Amp | pJN105 based expression vector containing LasR with A to G nucleotide substitution at position 664, resulting in T222A amino acid substitution | This work |
| pIY86/ pJN105LasR-C188A-Amp | pJN105 based expression vector containing LasR with TG to GC nucleotide substitution at position 562 and 563, resulting in C188A amino acid substitution | This work |

| | | |
|---|---|---|
| pIY87/ pJN105LasR-I190A-Amp | pJN105 based expression vector containing LasR with AT to GC nucleotide substitution at position 568 and 569, resulting in I190A amino acid substitution | This work |
| pIY88/ pJN105LasR-K192A-Amp | pJN105 based expression vector containing LasR with AA to GC nucleotide substitution at position 574 and 575, resulting in K192A amino acid substitution | This work |
| pIY89/ pJN105LasR-T193A-Amp | pJN105 based expression vector containing LasR with A to G nucleotide substitution at position 577, resulting in T193A amino acid substitution | This work |
| pIY90/ pJN105LasR-E196A-Amp | pJN105 based expression vector containing LasR with A to C nucleotide substitution at position 587, resulting in E196A amino acid substitution | This work |
| pIY91/ pJN105LasR-S198A-Amp | pJN105 based expression vector containing LasR with T to G nucleotide substitution at position 592, resulting in S198A amino acid substitution | This work |
| pIY92/ pJN105LasR-V199A-Amp | pJN105 based expression vector containing LasR with GG to CC nucleotide substitution at position 596 and 597, resulting in V199A amino acid substitution | This work |
| pIY93/ pJN105LasR-I200A-Amp | pJN105 based expression vector containing LasR with AT to GC nucleotide substitution at position 598 and 599, resulting in I200A amino acid substitution | This work |
| pIY94/ pJN105LasR-C201A-Amp | pJN105 based expression vector containing LasR with TG to GC nucleotide substitution at position 601 and 602, resulting in C201A amino acid substitution | This work |
| pIY95/ pJN105LasR-N202A-Amp | pJN105 based expression vector containing LasR with AA to GC nucleotide substitution at position 603 and 604, resulting in N202A amino acid substitution | This work |
| pIY96/ pJN105LasR-C203A-Amp | pJN105 based expression vector containing LasR with TG to GC nucleotide substitution at position 607 and 608, resulting in C203A amino acid substitution | This work |
| pIY97/ pJN105LasR-S204A-Amp | pJN105 based expression vector containing LasR with T to G nucleotide substitution at position 610, resulting in S204A amino acid substitution | This work |
| pIY98/ pJN105LasR-E205A-Amp | pJN105 based expression vector containing LasR with A to C nucleotide substitution at position 614, resulting in E205A amino acid substitution | This work |
| pIY99/ pJN105LasR-N207A-Amp | pJN105 based expression vector containing LasR with AA to GC nucleotide substitution at position 619 and 620, resulting in N207A amino acid substitution | This work |

| | | |
|---|---|---|
| pIY100/ pJN105LasR-N209A-Amp | pJN105 based expression vector containing LasR with AT to GC nucleotide substitution at position 625 and 626, resulting in N209A amino acid substitution | This work |
| pIY101/ pJN105LasR-M212A-Amp | pJN105 based expression vector containing LasR with AA to GC nucleotide substitution at position 634 and 635, resulting in M212A amino acid substitution | This work |
| pIY102/ pJN105LasR-G213A-Amp | pJN105 based expression vector containing LasR with G to C nucleotide substitution at position 638, resulting in G213A amino acid substitution | This work |
| pIY103/ pJN105LasR-I215A-Amp | pJN105 based expression vector containing LasR with AT to GC nucleotide substitution at position 643 and 644, resulting in I215A amino acid substitution | This work |
| pIY104/ pJN105LasR-R216A-Amp | pJN105 based expression vector containing LasR with CG to GC nucleotide substitution at position 646 and 647, resulting in R216A amino acid substitution | This work |
| pIY105/ pJN105LasR-R217A-Amp | pJN105 based expression vector containing LasR with CG to GC nucleotide substitution at position 649 and 650, resulting in R217A amino acid substitution | This work |
| pIY106/ pJN105LasR-F219A-Amp | pJN105 based expression vector containing LasR with TT to GC nucleotide substitution at position 655 and 656, resulting in F219A amino acid substitution | This work |
| pIY107/ pJN105LasR-G220A-Amp | pJN105 based expression vector containing LasR with G to C nucleotide substitution at position 659, resulting in G220A amino acid substitution | This work |
| pIY108/ pJN105LasR-V221A-Amp | pJN105 based expression vector containing LasR with T to C nucleotide substitution at position 662, resulting in V221A amino acid substitution | This work |
| pIY109/ pJN105LasR-R225A-Amp | pJN105 based expression vector containing LasR with CG to GC nucleotide substitution at position 673 and 674, resulting in R225A amino acid substitution | This work |
| pIY110/ pJN105LasR-V226A-Amp | pJN105 based expression vector containing LasR with T to C nucleotide substitution at position 677, resulting in V226A amino acid substitution | This work |
| pIY111/ pJN105LasR-I229A-Amp | pJN105 based expression vector containing LasR with AT to GC nucleotide substitution at position 685 and 686, resulting in I229A amino acid substitution | This work |
| pIY112/ pJN105LasR-M230A-Amp | pJN105 based expression vector containing LasR with AT to GC nucleotide substitution at position 688 and 689, resulting in M230A amino acid substitution | This work |

| | | |
|---|---|---|
| pIY113/ pJN105(Ap)-LasR-Ctrl | pJN105 based expression vector containing wild-type *lasR* | This work |
| pIY131/ pJN105(Ap)-RhlR-Ctrl | pJN105 based expression vector containing wild-type *rhlR* | This work |
| pIY134/ pJN105(Ap)LasR R216W | pJN105 based expression vector containing LasR with a C to T substitution at nucleotide 646 resulting in an R216W amino acid change | This work |
| pIY135/ pJN105(Ap)LasR R216W-M230I | pJN105 based expression vector containing LasR with a C to T substitution at nucleotide 646 and a G to A substitution at nucleotide 690 resulting in the dual R216W-M230I amino acid change | This work |
| pIY136/ pJN105(Ap)LasR A231V | pJN105 based expression vector containing LasR with a C to T substitution at nucleotide 692, resulting in an A231V amino acid change | This work |
| pIY137/ pJN105(Ap)LasR V226I | pJN105 based expression vector containing LasR PCR amplified from V226I natural variant strain (4) | This work |
| pEXG2-lasR_KO | pEXG2-based *lasR* deletion construct containing *lasR* gene and 500 bp up- and down-stream | (9) |
| pIY129/ pEXG2lasR_S179A_KI | pEXG2-based knock-in construct containing *lasR* with 500 bp up and downstream and the S179A variant described above | This work |
| pIY144/ pEXG2lasR_N202A_KI | pEXG2-based knock-in construct containing *lasR* with 500 bp up and downstream and the N202A variant described above | This work |
| pIY121/ pEXG2lasR_N207A_KI | pEXG2-based knock-in construct containing *lasR* with 500 bp up and downstream and the N207A variant described above | This work |
| pIY123/ pEXG2lasR_R217A_KI | pEXG2-based knock-in construct containing *lasR* with 500 bp up and downstream and the R217A variant described above | This work |
| pIY145/ pEXG2lasR_T222A_KI | pEXG2-based knock-in construct containing *lasR* with 500 bp up and downstream and the T222A variant described above | This work |
| pASD01 | pEXG2 containing *lasR* with 500 bp up and downstream and a C to T substitution at nucleotide 646, resulting in an R216W amino acid change in LasR | This work |
| pASD02 | pEXG2 containing *lasR* with 500 bp up and downstream and containing a C to T substitution at nucleotide 646 and a G to A substitution at nucleotide 690 resulting in the dual R216W - M230I amino acid change in LasR | This work |
| pASD03 | pEXG2 containing *lasR* with 500 bp up and downstream and a C to T substitution at nucleotide 692, resulting in an A231V amino acid change in LasR | This work |

| | | |
|---|---|---|
| pUC18-miniTN7TGm-gfp | used for the integration of *gfp* at the *att* site | (10) |
| pUC18-miniTN7TGm-mCherry | Derived from pUC18-mini-Tn7T-Gm; used for the integration of mCherry at the *att* site | (11) |
| pFlp2 | Used to express the *flp* recombinase gene for excising a resistance marker from the *mCherry* cassette delivered in the pUC18-mini-Tn7TGm-mCherry | (10) |

**References**

1. Jacobs MA, Alwood A, Thaipisuttikul I, Spencer D, Haugen E, Ernst S, Will O, Kaul R, Raymond C, Levy R, Chun-Rong L, Guenthner D, Bovee D, Olson MV, Manoil C. 2003. Comprehensive transposon mutant library of *Pseudomonas aeruginosa*. Proc Natl Acad Sci USA 100:14339–14344.

2. Wang M, Schaefer AL, Dandekar AA, Greenberg EP. 2015. Quorum sensing and policing of *Pseudomonas aeruginosa* social cheaters. Proc Natl Acad Sci USA 112:2187–2191.

3. Siehnel R, Traxler B, An DD, Parsek MR, Schaefer AL, Singh PK. 2010. A unique regulator controls the activation threshold of quorum-regulated genes in *Pseudomonas aeruginosa*. Proc Natl Acad Sci USA 107:7916–7921.

4. Bellamoroso KR, Kostylev M, Smalley NE, Greenberg EP, Dandekar AA. 2026. Inactivation of MexT in *Pseudomonas aeruginosa* PAO1 destabilizes cooperation and favors the emergence of a unique quorum sensing variant. J Bacteriol 208:e00434-25.

5. Tseng BS, Zhang W, Harrison JJ, Quach TP, Song JL, Penterman J, Singh PK, Chopp DL, Packman AI, Parsek MR. 2013. The extracellular matrix protects *Pseudomonas aeruginosa* biofilms by limiting the penetration of tobramycin. Environmental Microbiology 15:2865–2878.

6. Kostylev M, Kim DY, Smalley NE, Salukhe I, Greenberg EP, Dandekar AA. 2019. Evolution of the *Pseudomonas aeruginosa* quorum-sensing hierarchy. Proc Natl Acad Sci USA 116:7027–7032.

7. Simon R, Priefer U, Pühler A. 1983. A broad host range mobilization system for in vivo genetic engineering: transposon mutagenesis in Gram negative bacteria. Nat Biotechnol 1:784–791.

8. Zheng X, Gomez-Rivas EJ, Lamont SI, Daneshjoo K, Shieh A, Wozniak DJ, Parsek MR. 2024. The surface interface and swimming motility influence surface-sensing responses in *Pseudomonas aeruginosa*. Proc Natl Acad Sci USA 121:e2411981121.

9. Wellington Miranda S, Cong Q, Schaefer AL, MacLeod EK, Zimenko A, Baker D, Greenberg EP. 2021. A covariation analysis reveals elements of selectivity in quorum sensing systems. eLife 10:e69169.

10. Choi K-H, Gaynor JB, White KG, Lopez C, Bosio CM, Karkhoff-Schweizer RR, Schweizer HP. 2005. A Tn7-based broad-range bacterial cloning and expression system. Nat Methods 2:443–448.

11. Zhao K, Tseng BS, Beckerman B, Jin F, Gibiansky ML, Harrison JJ, Luijten E, Parsek MR, Wong GCL. 2013. Psl trails guide exploration and microcolony formation in *Pseudomonas aeruginosa* biofilms. Nature 497:388–391.
